# Low-frequency motor cortex EEG predicts four levels of rate of change of force during ankle dorsiflexion

**DOI:** 10.1101/2022.11.02.514949

**Authors:** Rory O’Keeffe, Seyed Yahya Shirazi, Alessandro Del Vecchio, Jaime Ibáñez, Natalie Mrachacz-Kersting, Ramin Bighamian, JohnRoss Rizzo, Dario Farina, S. Farokh Atashzar

## Abstract

The movement-related cortical potential (MRCP) is a low-frequency component of the electroencephalography (EEG) signal recorded from the motor cortex and its neighboring cortical areas. Since the MRCP encodes motor intention and execution, it may be utilized as an interface between patients and neurorehabilitation technologies. This study investigates the EEG signal recorded from the Cz electrode to discriminate between four levels of rate of force development (RFD) of the tibialis anterior muscle. For classification, three feature sets were evaluated to describe the EEG traces. These were (i) *MRCP morphological characteristics* in the *δ*-band such as amplitude and timing, (ii) *MRCP statistical characteristics* in the *δ*-band such as mean, standard deviation, and kurtosis, and (iii) *wideband time-frequency features* in the 0.5-90 Hz range. Using a support vector machine for classification, the four levels of RFD were classified with a mean (SD) accuracy of 82% (7%) accuracy when using the time-frequency feature space, and with an accuracy of 75% (12%) when using the MRCP statistical characteristics. It was also observed that some of the key features from the statistical and morphological sets responded monotonically to the intensity of the RFD. Examples are slope and standard deviation in the (0, 1)s window for the statistical, and *min*_1_ and *min_n_* for the morphological sets. This monotonical response of features explains the observed performance of the *δ*-band MRCP and corresponding high discriminative power. Results from temporal analysis considering the pre-movement phase ((-3, 0)s) and three windows of the post-movement phase ((0, 1)s, (1, 2)s, and (2, 3)s)) suggest that the complete MRCP waveform represents high information content regarding the planning, execution, duration, and ending of the isometric dorsiflexion task using the tibialis anterior muscle. Results shed light on the role of *δ*-band in translating to motor command, with potential applications in neural engineering systems.

## Introduction

Motor-related potentials are the aggregation of neural activity in the motor cortex. The neural activity can be related to sensory integration, motor preparation, motor learning, and execution of a motor task [1–8]. The dorsal premotor cortex (PMd) is believed to have a central role in motor planning [9–11]. The supplementary motor area (SMA) is known to act as a hub for the integration of cognitive, motor and sensory information [12, 13]. The summations of neuron-level activities generate a local field potential (LFP), which is believed to be the source of the electrical signals recorded at the scalp, known as electroencephalography (EEG) [14–16]. An EEG recording from the central region of the brain (especially near the Cz electrode location) represents the LFPs generated by the motor-related potentials at the premotor cortex, SMA and the medial areas of the motor cortex [17–20].

Motor-related potentials can discriminate motor impairments, quantify a variety of tasks and also monitor the improvements after rehabilitation, especially for stroke and other central nervous system impairments [3, 21–24]. Furthermore, cognitive impairments such as attention deficit hyperactivity disorders can degrade the sensorimotor processing of the brain, which specifically reflects on the SMA and motor cortex [25]. Motor-related potentials are especially important from the brain-computer interface perspective because they provide information about the sensorimotor processing which precedes motor actions [21, 26, 27]. For cue-based movements, the PMd is believed to have a major role in this pre-movement activity which can be captured as negative deflections in the motor-related potentials [28, 29].

The negative deflections in the neural trace are the most prominent features of the movement-related cortical potential (MRCP). MRCP is a low-frequency (0-4 Hz, *δ*-band) EEG signal component generated in response to the planning and execution of a cued or self-paced voluntary movement [30–34]. For lower-limb activity, the MRCP is usually recorded by EEG electrodes at the mid-central area of the brain, such as Cz, which predominantly reflects sources from the SMA [35, 36], premotor and primary motor cortices [4,37]. The MRCP consists of three components, called the readiness potential (RP), motor potential, and movement-monitoring potential (MMP), corresponding to movement planning, execution, and control, respectively [38,39]. The RP is also called a Bereitschafts potential in the context of self-paced movements and is a negative deflection that may begin as early as 2s prior to the movement onset [38,40]. The RP consists of two phases, the second of which usually has a steeper slope and maximum amplitude over the primary motor cortex [40]. Both the RP and MMP have been shown to be related to the task’s kinetic parameters, for example, force and rate of force development (RFD) [31, 38]. An imagined MRCP is also produced in response to motor imagery and usually has lower amplitude than an executed movement [38]. The MRCP has been used to predict and understand the execution of tasks in healthy adults and patients with Amyotrophic Lateral Sclerosis, tremor, Parkinson’s disease, and stroke [34, 39, 41, 42].

Mapping the MRCP to multiple distinct levels of a single kinetic parameter is an important next step for brain-computer interface (BCI) and neurorehabilitation technologies, yet efforts to date have struggled to attain high classification accuracy with >2 levels of a single parameter [43, 44]. It has been demonstrated that the detection of an MRCP can be reliably performed with low latency (high temporal resolution) [45, 46]. Additionally, the MRCP can distinguish left vs. right hand, and foot movements [47, 48]. When using the MRCP to map to a kinetic parameter, distinguishing two levels of force or RFD has been successful (accuracy > 74% for force, accuracy > 81% for RFD) [49]. Successful classification of multiple RFD levels is particularly important as the BCI can aid the user in performing the task at the intended speed. This provides an additional motor dimension beyond the start and stop of the task and may be crucial for neurorehabilitation technologies. This study investigates the potential discrimination of four RFD levels with MRCPs, comparing features both in the *δ*-band and in the full EEG band. Moreover, we performed a detailed analysis of the MRCP’s characteristics, with a focus on the discriminative power of features from the *δ*-band at Cz.

Our first hypothesis is that four levels of RFD can be accurately discriminated using full-band EEG features from a larger scalp area spanning nine central electrodes. Secondly, we hypothesize that the MRCP’s temporal *δ*-band features from the Cz electrode can achieve a comparable classification accuracy.

## Materials and Methods

Five healthy volunteers (males, aged 20-28 years), without any prior BCI experience participated in the study. All subjects gave their signed consent to the study, which was conducted in accordance with the principles outlined in the Helsinki Declaration and approved by the University College London Department of Clinical and Movement Neurosciences (approval date: April 26th 2017, approval reference number: 10037/001).

### Experimental Protocol

Each subject was seated in a chair with their leg constrained (NEG1, OT Bioelettronica, Torino, Italy) and ankle fixed to a pedal with an attached force transducer (Fig. 1(a)). Subjects were instructed to perform a defined rate of isometric dorsiflexion of the dominant-sided (right in all subjects) ankle. Four rates of force development (RFD) were defined based on the time interval to reach a target force level of 60% of maximal voluntary contraction (MVC). With real-time visual feedback, subjects followed a triangular profile (Fig. 1(b)) from 0 to 60% MVC with the durations of: (i) 3s, (ii) 2s, (iii) 1s and (iv) 0.5s, corresponding to RFDs of (i) 20% MVC/s (Slow), (ii) 30% MVC/s (Medium), (iii) 60% MVC/s (Fast), and (iv) 120% MVC/s (Ballistic). Each experimental session started with the measurement of the maximum voluntary contraction (MVC) force, so that the RFD levels could be defined. A training phase (approximately 5 min) was included to let the subjects familiarize themselves with executing the defined force profile while receiving visual feedback. The subjects were guided to execute a set of 25 isometric ankle dorsiflexion repetitions at a certain RFD, interspersed with 5s of resting periods. Between each set, a break of 5 min was provided.

**Figure 1:**
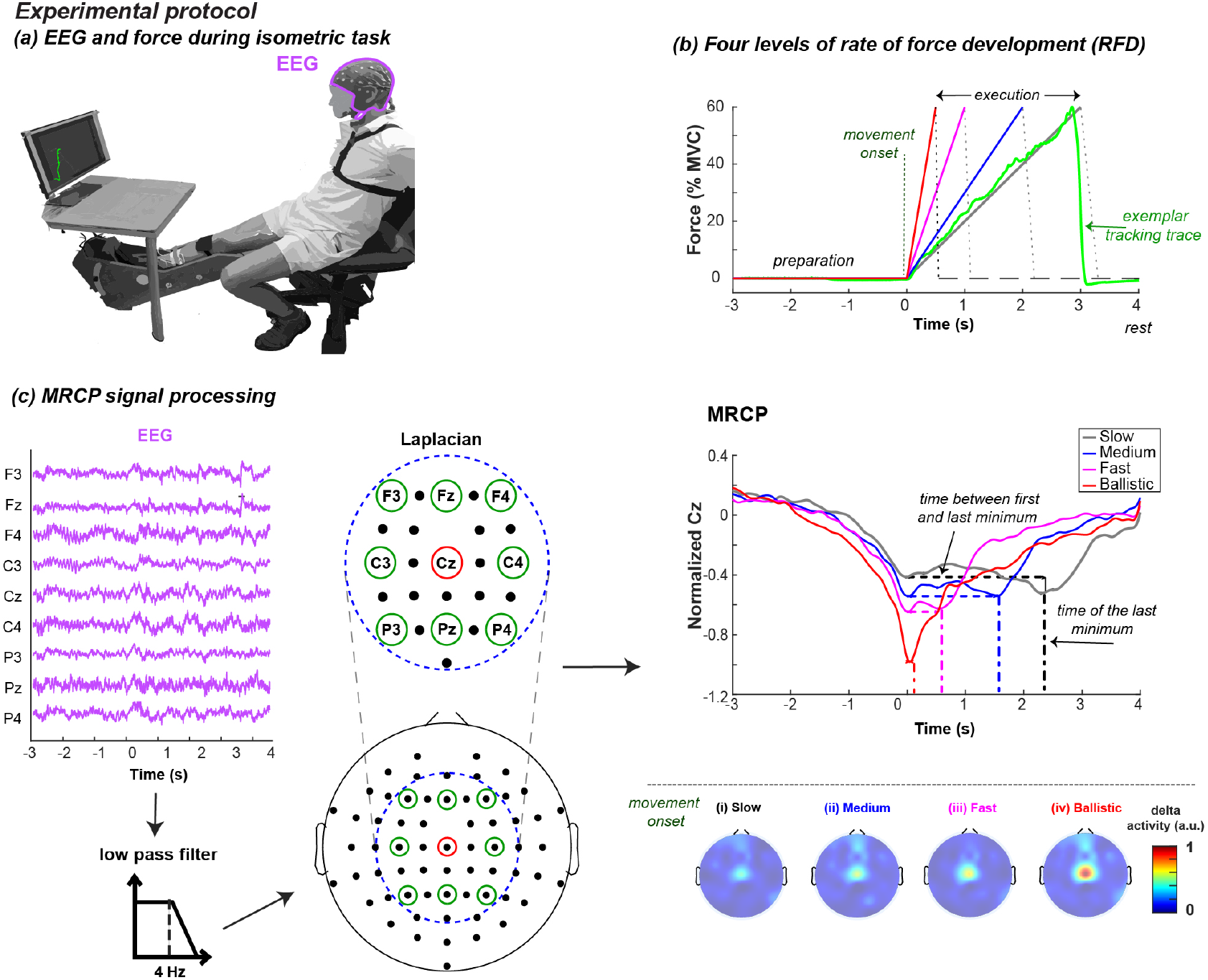
**(a)** EEG signals were recorded using a 10-10 system and force signals were measured using a load cell while subjects performed guided isometric ankle dorsiflexion. **(b)** Subjects reached 60% maximum voluntary contraction (MVC) with four durations (0.5s, 1s, 2s and 3s), giving four levels of rate of change of force (Ballistic: 120% MVC/s, Fast: 60% MVC/s, Medium: 30% MVC/s and Slow: 20% MVC/s). **(c)** The MRCP is produced by performing a Laplacian spatial filter with eight electrodes surrounding Cz and a low pass filter at 4 Hz. Several MRCP morphological characteristics, such as the time features of the the minima seem to scale with the task duration. Heat maps indicate that *δ*-band activity is highest for Cz and scales in proportion to RFD.

### Data Acquisition

EEG data were wirelessly recorded from 64 electrodes assembled on an active-electrode cap (BrainAmp, Brain Products GmbH, Germany) joined with a Ses-santaquattro transmitter module (OTBioelettronica, Turin, Italy) at 2048 Hz. The force signal was recorded from a force transducer mounted on a pedal and connected to an amplifier (Quattrocento, OTBioelettronica, Turin, Italy). At the start of each set of isometric dorsiflexions, a trigger pulse was sent to the EEG recording system when the force signal recording started, such that the EEG and force signals could be synchronized later. The movement onset was defined as the time at which the force signal reached 10% of its peak trial value (approximately 6% MVC, as in Fig. 1(b)). Using this movement onset time as 0s for each trial, a [-3, 4] s window was used to investigate the EEG signals for each trial.

### EEG Processing

Nine EEG electrodes of interest, including and neighboring Cz, i.e., F3, Fz, F4, C3, Cz, C4, P3, Pz and P4, were used for the analysis (Fig. 1(c)). EEG signals from these electrodes were filtered with (i) a zero-phase, 2nd-order Butterworth high pass filter with a cutoff frequency of 0.5 Hz, (ii) a zero-phase, 4th-order Butterworth band stop filter (47.5-52.5 Hz) to remove the line noise and (iii) a zero-phase, 4th-order Butterworth low pass filter with a cut off frequency of 90 Hz.

### MRCP Analysis

To extract the MRCP, first a Laplacian spatial filter was applied to the Cz electrode to enhance the spatial resolution (Fig. 1(c)). The signal was then low-pass filtered at 4 Hz with a 4th-order zero-phase Butterworth filter. The mean MRCP, *μ_sam_,* across all trials of all subjects was computed, for each RFD, and the results can be seen in Fig. 2(a) and Fig. 2(b). Additionally, the sample standard deviation, 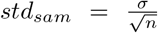, across all trials of all subjects was computed and used to compute the MRCP’s 95% confidence interval *(CI_95_,* Fig. 2(a)) as *CI*_95_ = *μ_sam_* ± (*Z*_95_)(*std_sam_*)= *μ_sam_* ± (1.96)(std_sam_). The *δ*-band activity was plotted, resulting in the activation heat maps as explained in the following. After bandpass filtering, each of the 64 recorded electrodes was spatially filtered with an appropriate Laplacian configuration. The configuration was chosen based on the position of the electrode of interest, and its surrounding electrodes [50]. The median magnitude of each electrode’s time domain signal across all subject trials was computed, and the median in a 100 ms window produced a singular value for the topographical heat maps (Fig. 2(c)). The 100 ms window was centered at the movement onset for the maps in the top row (centered at 0s for all classes). For the bottom row, the window was centered at the planned movement end (centered at 3s for Slow, at 2s for Medium, at 1s for Fast and at 0.5s for Ballistic). Each electrode’s value was normalized using the maximum across the eight plots in Fig. 2(c).

**Figure 2:**
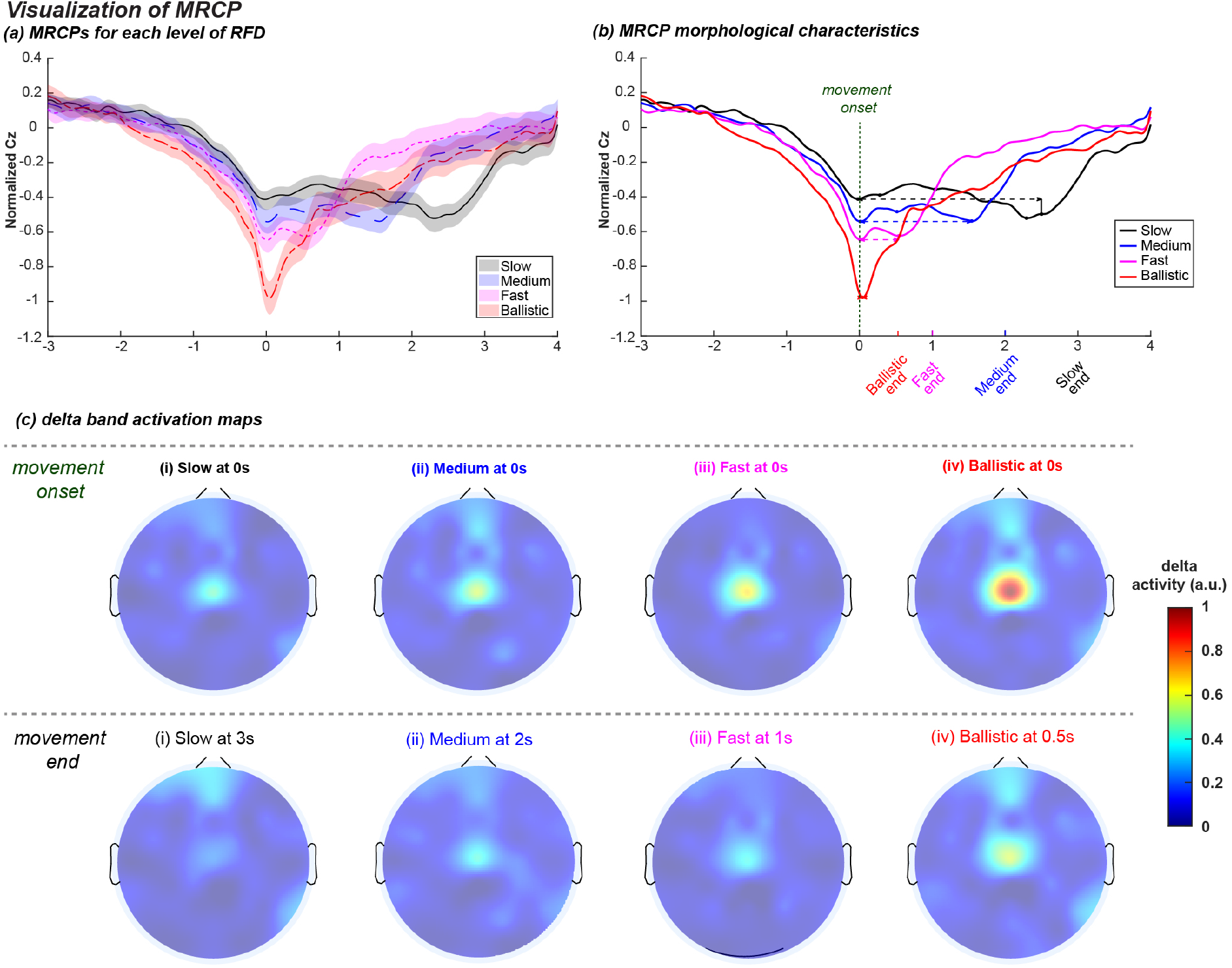
**(a)** Mean (center line) MRCP with 95% confidence interval (shaded area) across all subjects. **(b)** Mean MRCP for each class of RFD, where the dots signify local minima. The time between the first and last local minimum (denoted by a dashed line) is closely related to the duration of the force application. The slope of the late readiness potential (RP2 slope) for each level of RFD is displayed. **(c)** Topographical plots of *δ*-band activation for the recorded electrodes. The maps in the top row considered the movement onset (0s), while the maps in the bottom row considered the planned movement end as the time window center. The dominant *δ*-band activation appears to be at Cz and is monotonically ascending with the level of RFD (with Slow having the smallest and Ballistic having the largest). The activation of Cz trended higher at the movement onset than at the end. For each subject and trial, the median activity was quantified for the 100ms window around the specified time. Here, the maps represent the median across all subject trials and were normalized to the maximum activity across the eight plots.

### Feature Set 1: MRCP Morphological Characteristics

The following features were considered as the morphological characteristics of the MRCP at Cz.

i. The slope of the late readiness potential (RP2), i.e., the slope of the MRCP between −0.5s and 0s
ii. The first local minimum’s value, *min*_1_, traditionally called peak negativity (PN)
iii. The time at which *min*_1_ occurred, *t*_*min*_1__
iv. The number of local minima, *N_min_*
v. The last local minimum’s value, *min_n_*
vi. The time at which *min_n_* occurred, *t_minn_*.

### Feature Set 2: MRCP Statistical Characteristics

The following were considered as the statistical characteristics of the MRCP at Cz in the temporal domain:

i. Mean value
ii. Standard deviation
iii. Mean absolute value
iv. Trapezoidal integral (area under the curve)
v. Slope between the initial and final value
vi. Slope sign change (SSC) count
vii. Mean 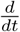 of MRCP signal
viii. Skewness *s*, which gives a measurement of how asymmetrically a signal is distributed about the mean, can be computed as

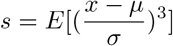
ix. Kurtosis *k,* which gives a positive measurement if a signal has few outliers, and a negative measurement when there are many, can be computed as:

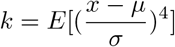
x. Shannon’s entropy

The above features were computed for seven time windows, from 3s before the movement onset to 4s after the movement onset (each time window is one second long). Features from all time windows were pooled to train and evaluate the performance of the classifier (Fig. 5). We also evaluated the classifier performance when it was only trained on individual time windows (e.g., 1 to 2 sec, or 3 to 4 sec), which is depicted in Fig. 6 and Table 1.

**Figure 3:**
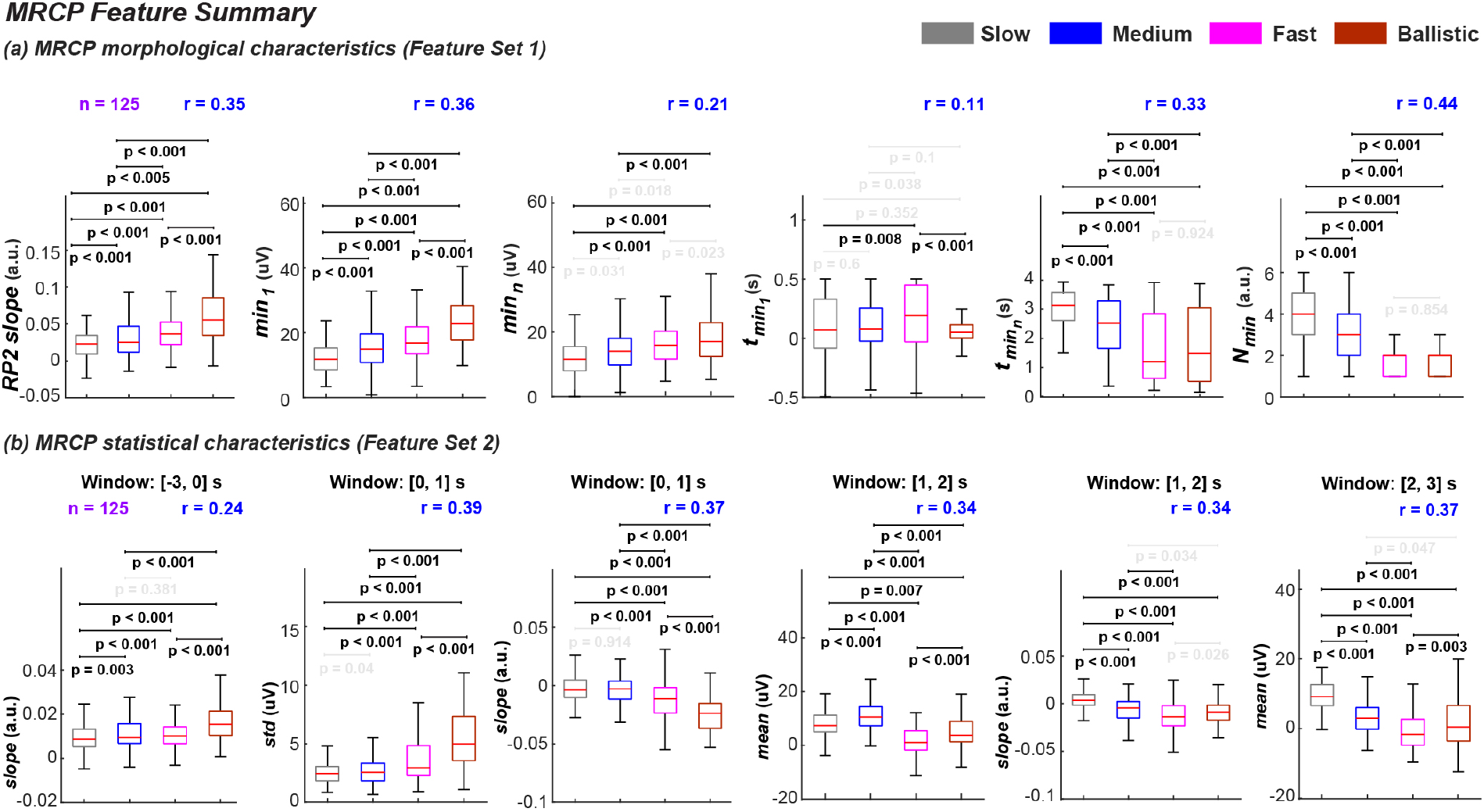
Statistical results of MRCP features. The features were computed for all trials (*n* = 125). Post-hoc adjusted significance level for all features: *α*’= 0.083. **(a)** Results for MRCP morphological characteristics. *RP*2 slope and *min*_1_ monotonically ascended with the level of RFD. *t_minn_* and *N_min_* showed monotonical descension from Slow to Fast. The rank biserial correlation of |*r_rb_*| = 0.44 for *N_min_* indicated the largest effect size in the feature set. **(b)** The statistical analysis for each of six exemplar MRCP attributes is shown. In the [0,1] s time window, the standard deviation monotonically ascended while the slope monotonically descended from Medium to Ballistic. The slope in the [1,2] s window and the mean in the [2,3] s window showed monotonical descension from Slow to Fast. The largest effect size |*r_rb_*| = 0.39 was observed for the standard deviation in the [0,1] s window.

**Figure 4:**
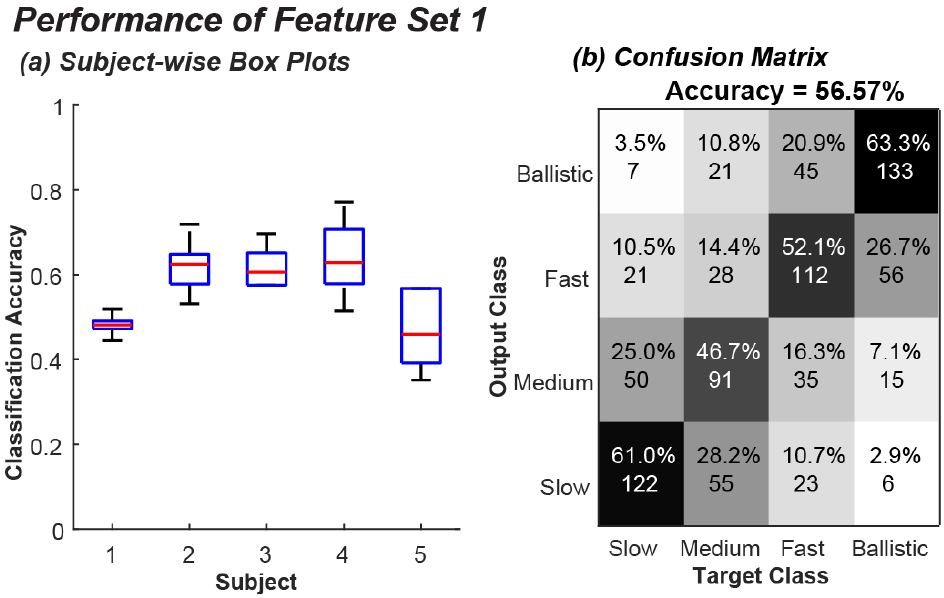
Classification accuracy from MRCP morphological characteristics (Feature Set 1). **(a)** Subject-wise box plots of classification accuracy. The distribution of classification accuracy across all trials of the Monte Carlo repetitions for each subject is shown. **(b)** Overall confusion matrix. The classification results from all subjects are pooled and the mean accuracy is shown above.

**Figure 5:**
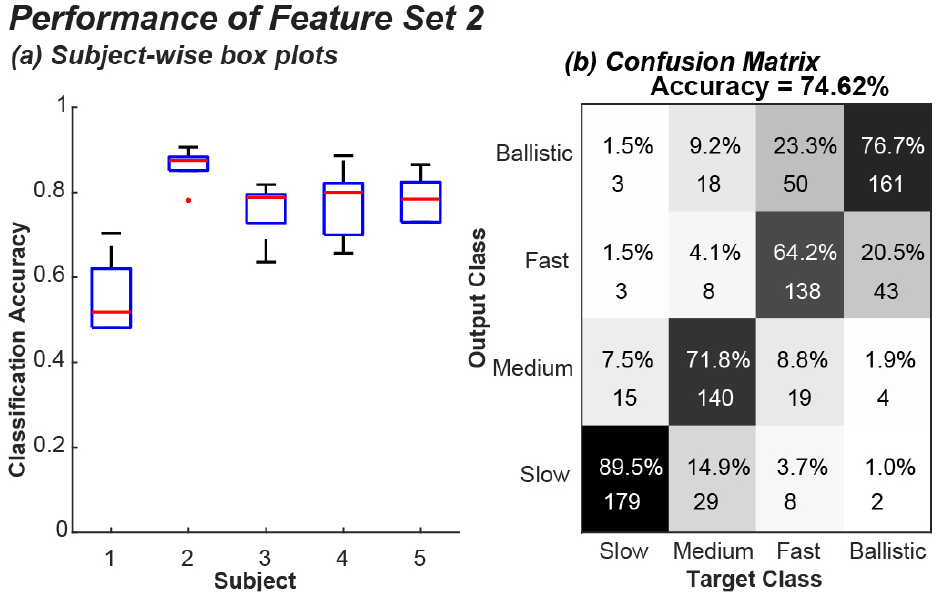
Classification accuracy from MRCP statistical characteristics (Feature Set 2) from all time windows. **(a)** Subject-wise box plots of classification accuracy. The distribution of classification accuracy across all trials of the Monte Carlo repetitions for each subject is shown. **(b)** Overall confusion matrix. The classification results from all subjects are pooled and the mean accuracy is shown above.

**Figure 6:**
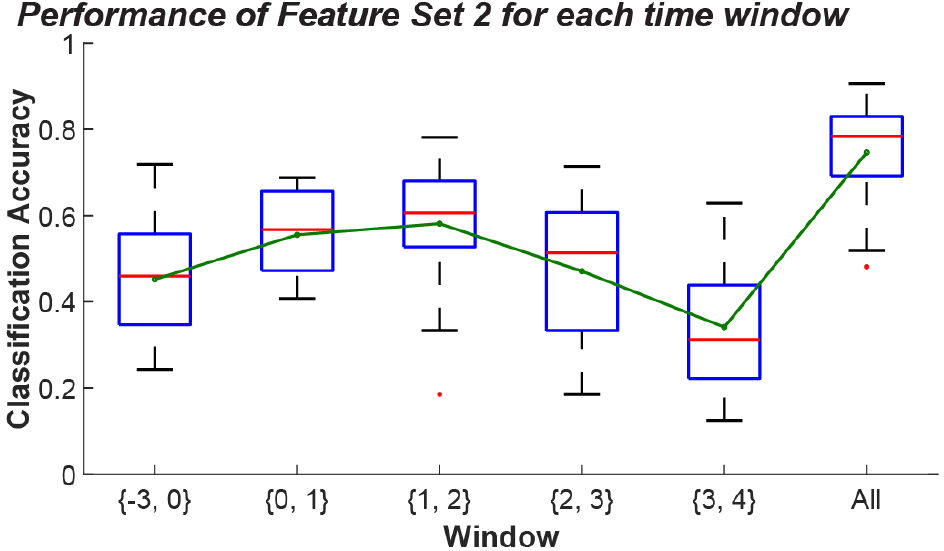
Window-wise classification accuracy from statistical characteristics (Feature Set 2). Each of the first five box plots depicts the accuracy obtained when classifying the level of rate of change of force with features across just that time window. The final box plot shows the accuracy when classifying with features pooled from all time windows. Each box plot shows a *subjects × trials* distribution for the given time window. The green line plot indicates the mean classification accuracy.

**Table 1:**
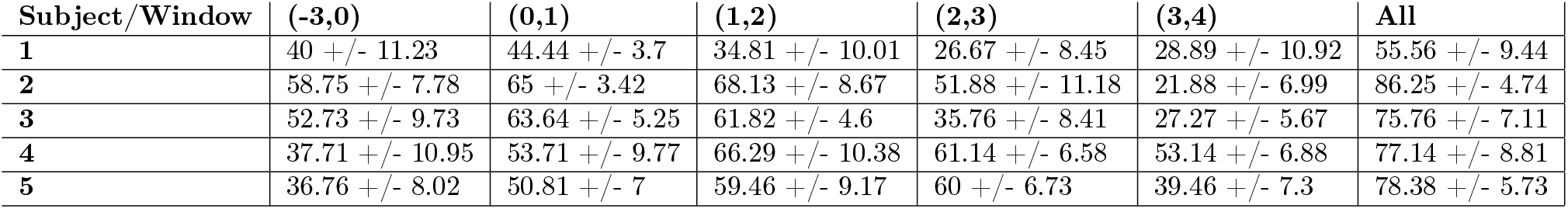
Window-wise classification accuracy (CA) using temporal MRCP features. The *mean* ± *std* CA across all trials of the Monte Carlo repetitions for each subject is shown.

| Subject/Window | (-3,0) | (0,1) | (1,2) | (2,3) | (3,4) | All |
| --- | --- | --- | --- | --- | --- | --- |
| 1 | 40 +/- 11.23 | 44.44 +/- 3.7 | 34.81 +/- 10.01 | 26.67 +/- 8.45 | 28.89 +/- 10.92 | 55.56 +/- 9.44 |
| 2 | 58.75 +/- 7.78 | 65 +/- 3.42 | 68.13 +/- 8.67 | 51.88 +/- 11.18 | 21.88 +/- 6.99 | 86.25 +/- 4.74 |
| 3 | 52.73 +/- 9.73 | 63.64 +/- 5.25 | 61.82 +/- 4.6 | 35.76 +/- 8.41 | 27.27 +/- 5.67 | 75.76 +/- 7.11 |
| 4 | 37.71 +/- 10.95 | 53.71 +/- 9.77 | 66.29 +/- 10.38 | 61.14 +/- 6.58 | 53.14 +/- 6.88 | 77.14 +/- 8.81 |
| 5 | 36.76 +/- 8.02 | 50.81 +/- 7 | 59.46 +/- 9.17 | 60 +/- 6.73 | 39.46 +/- 7.3 | 78.38 +/- 5.73 |

### Feature Set 3: Wideband Time-frequency Features

The short-time Fourier transform (STFT) was performed to generate the benchmarking feature space on the nine electrodes of interest (F3, Fz, F4, C3, Cz, C4, P3, Pz and P4), with 50% overlapping Hamming windows of 2s. The nine-electrode STFT formed the spectrotemporal feature space in the broad frequency range of interest (0.5-90 Hz).

### Statistical Analysis

The relationship of features with respect to the four levels of RFD was analyzed for the MRCP characteristics (Fig. 3). Each feature distribution consisted of the computed values for all trials (*n* = 125). The Kolmogorov-Smirnov test for normality rejected the normal distribution hypothesis for all distributions; therefore, non-parametric statistical tests were used in our analysis. The Friedman test was used to compare levels of RFD. The Wilcoxon signed-rank test was used as a post-hoc test if the Friedman test revealed significance. The significance level, *α*, for all tests was initially set at 0.05. The Bonferroni correction was applied to adjust for multiple comparisons, dividing *α* by the number of comparisons such that the adjusted significance level was *α*’ = 0.05/6 = 0.0833. The effect size of the non-normal distributions was quantified by using the rank-biserial correlation [51]. In this regard, |*r_rb_*| was used for measuring the rank-biserial correlation. A higher value means that the effect size is larger.

### Machine Learning Classification

For all three feature sets, a Support Vector Machine (SVM) classifier with a linear kernel function was used. For a given subject, there were ~25 trials of each RFD class (Slow, Medium, Fast, Ballistic). A training and test set was defined for each subject. The test portion of the data was 1/3 of the trials, but rather than using conventional 3-fold cross-validation, the more robust Monte Carlo method was used [52]. On each repetition of the Monte Carlo method, the test set is randomly selected from the data, and the training set constitutes the remaining samples. The model is trained before computing the accuracy of the trained model on the test set. Five repetitions of training and testing were used for all results shown to compute the accuracy.

## Results

### Feature Set 1: MRCP Morphological Characteristics

Increases in RFD resulted in identifiable changes in the MRCP signal features. The *CI_95_* plots demonstrated significant differences between the group average MRCPs across the RFDs (Fig. 2(a)). The value of the first local minimum, the distance between the first and last local minima, and the MRCP slope before the execution, i.e., the readiness potential (RP2), seemed to be among the features contributing to the separation of the MRCPs and their respective *CI*_95_. Furthermore, the distance between the first and last MRCP minima seemed to scale with the force development time for each group (Fig. 2(a), Fig. 2(b)). The activation maps after Laplacian filter indicated monotonically increasing activity around the Cz area from the Slow, to Medium, to Fast, and to Ballistic RFDs. This activity tended to diminish at the end of the force development (Fig. 2(c)). For reference, the force development time is 3 s for the Slow group, 2 s for the Medium group, 1 s for the Fast group, and 0.5 s for the Ballistic group. The brain activity centered around Cz trended highest at the time points considered for Ballistic, particularly at the movement onset (Fig. 2(c)).

It should be noted that MRCP morphological characteristics showed significant differences depending on the level of RFD for pre-movement, movement onset and post-movement features (Fig. 3(a)). *RP*2 slope and *min*_1_ monotonically ascended with the level of RFD and all levels were statistically different from each other (*Friedman* 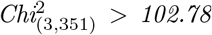, *p* < 0.05, post-hoc Wilcoxon signed-rank test: all six pairwise comparisons: *p* < 0.006). Features with a postmovement component such as *t_min_n__* and *N_min_* showed monotonical descension from Slow to Fast (*Friedman* 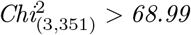, *p < 0.05, post-hoc Wilcoxon signed-rank test: 5/6 pairwise comparisons: p* < 0.001). The rank biserial correlation of |*r_rb_*| = 0.44 for *N_min_* indicated the largest effect size in the feature set. Five of the six features shown demonstrated a difference between Slow and Ballistic RFD (*post-hoc Wilcoxon signed-rank test: p* < 0.001).

Using the MRCP morphological characteristics as the feature set for the SVM algorithm demonstrated medium discriminative power in relation to the four levels of RFD (>50%), which is double the chance level for four classes (Fig. 4). The average overall accuracy was 57% across subjects and classes (Fig. 4). The confusion matrix for this feature set also suggested a strong discriminative power if only two RFD groups were considered (Fig. 4(b)). The cumulative accuracy of the adjacent RFD groups (i.e., Ballistic and Fast vs. Medium and Slow) is consistently at 80%. It should be noted that this feature set had only *six features* derived from the *δ*-band MRCP and robustly classified dual-level RFDs and performed well beyond the chance level for the four-level RFD classification using SVM.

### Feature Set 2: MRCP Statistical Characteristics

MRCP statistical characteristics showed significant differences depending on the level of RFD, especially for post-movement features. The statistical analysis for six exemplar MRCP attributes is shown in Fig. 3(b). In the [0,1] s time window, the standard deviation showed monotonical ascension while the slope showed monotonical descension from Medium to Ballistic (*Friedman* 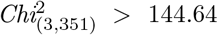, *p* < 0.05, *post-hoc Wilcoxon signed-rank test: 5/6 pairwise comparisons: p* < 0.001). The slope in the [1,2] s window and the mean in the [2,3] s window showed monotonical descension from Slow to Fast (*Friedman* 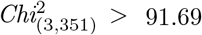, *p* < 0.05, *post-hoc Wilcoxon signed-rank test: 5/6 pairwise comparisons: p* < 0.004). The largest effect size |*r_rb_*| = 0.39 was observed for the standard deviation in the [0,1] s window. All six features shown demonstrated a difference between Slow and Ballistic RFD *(post-hoc Wilcoxon signed-rank test: p* < 0.008).

Using the ten MRCP statistical characteristics in Feature Set 2 provided RFD classification accuracy of ~ 75% (Fig. 5). Four out of the five subjects had >75% classification accuracy (Fig. 5(a), Table 1). The mean accuracy across subjects and RFD classes was 75% (Fig. 5(b)), with the highest discriminative accuracy at 89% for the Slow RFD. The cumulative accuracy of the two-level RFD with adjacent classes (i.e., Ballistic and Fast vs. Medium and Slow) reached 90% (Fig. 5(b)). Considering the MRCP statistical characteristics only from selected time windows as the SVM feature set reveals that the SVM classifier is likely most responsive to attributes in the [1,2] s window with accuracy ~ 60% (Fig. 6). The least discriminative power was for the [3,4] s time window with ~ 30% (around chance) accuracy. Interestingly, the median accuracy with all the time windows included was ~ 80%, which was greater than each individual time window (Fig. 6). The detailed per subject and window accuracy results also confirmed that for all subjects, the [0,1] s and [1,2] s windows provided the highest accuracies for each subject (Table 1).

### Feature Set 3: Wideband Time-frequency Features

The SVM classification using STFT full temporal and spectral feature set yielded >80% classification accuracy for the four levels of RFD. The subject-wise classification accuracy revealed >80% accuracy for four out of the five subjects (Fig. 7(a)), while the overall group accuracy for the STFT feature set was also at ~ 84% (Fig. 7(b)). Adjacent classes with the fastest RFDs (i.e., Fast and Ballistic) had a high mutual misclassification rate compared to the misclassification rate when the inter-class distance increased.

**Figure 7:**
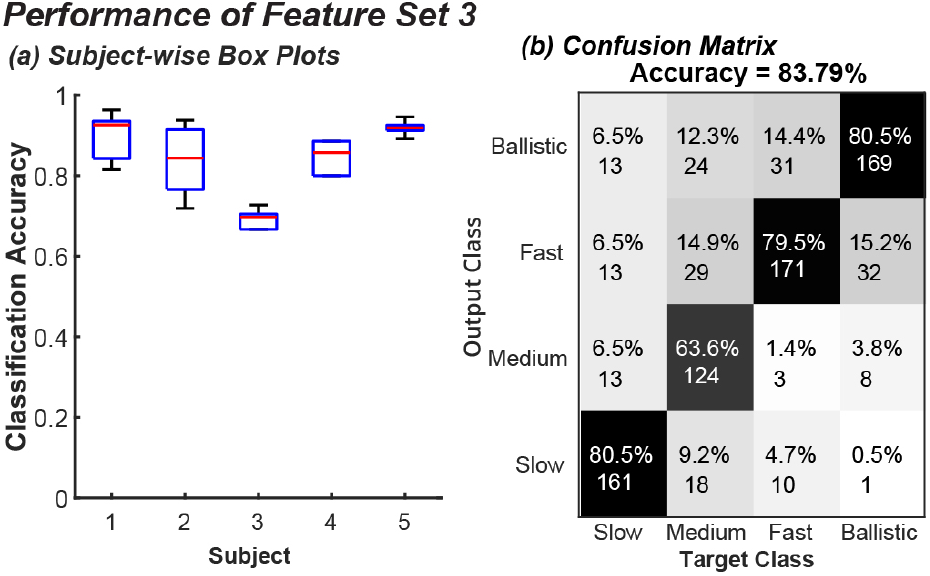
Classification accuracy from wideband STFT features (Feature Set 3) derived from the full spectrum of nine electrodes. **(a)** Subject-wise box plots of classification accuracy. The distribution of classification accuracy across all trials of the Monte Carlo repetitions for each subject is shown. **(b)** Overall confusion matrix. The classification results from all subjects are pooled and the mean accuracy is shown above.

## Discussion

Four RFD levels could be discriminated with ~ 75% accuracy when only using *δ*-band MRCP statistical characteristics. The complete MRCP waveform investigated for the first time in this paper for four levels of RFD showed high information content regarding the planning, execution, duration, and ending of the isometric dorsiflexion task using the tibialis anterior muscle. In this regard, the results showed that morphological features, such as the time between the first and last minima encapsulate critical information with potential use in BCI. We demonstrated that both MRCP morphological and statistical features present monotonic trends with respect to the RFD levels (Slow RFD was differentiated from Ballistic RFD in 11 out of 12 features shown, *p* < 0.001). This paper also shows that morphological features such as the number of minima *(N_min_)* and the time of the last minimum (*t_min_n__*), and statistical features such as MRCP mean and standard deviation (*std*), have as strong an effect size as previously explored features, such as RP2 slope and the magnitude of the first minimum (*min*_1_). Using the MRCP feature sets provides a transparent and interpretable SVM classifier that is not a “black box” and can inform researchers about the EEG waveform dynamics. The novel MRCP features are neurophysiological indications of the underlying mechanism of motor monitoring and execution by the motor cortex and the neighboring areas. This is the first study with successful four levels of RFD classification based on the MRCP. Our results illustrate the exciting possibility of MRCP to RFD mapping, which has strong potential applications, particularly in neurorehabilitation.

While the importance of some MRCP features, such as RP2 slope and *min*_1_ was explored previously [35, 53–56], we found that other features like the number of minima (N*min*) and the timing of the last minimum *(t_min_n__*) can be equally, if not more discriminative than the previously explored features (Fig. 3). We observed that *min_1_, min_n_,* and RP2 slope scaled in inverse while *N_min_* and *t_min_n__* scaled in direct proportion to the RFD levels (Figs. 2, 3). Peak negativity (PN, denoted here as *min*_1_) has been previously reported as the largest deflection in response to the movement intention [56, 57]. The scaling of the PN (*min*_1_) may be an indication of the level of error processing and corrective actions in the motor and premotor cortices [19,58,59]. Interestingly, we observed that the PN is not necessarily the most negative value reached over the task duration (e.g., Slow in Fig. 2(a)). Rather, the PN is just the first of a set of local minima,indicating that the error processing and movement monitoring can continue beyond the initial recruitment of the muscle. Based on the duration of the task, there might be even greater negativities following the *min*_1_. Previous work highlighted that two levels of RFD would produce two distinct *min*_1_’s [43]. Here, we show that four distinct levels of RFD provide four distinct *min*_1_’s (post-hoc *p* < 0.001 for all comparisons of in Fig. 3(a)). Our results support the view that the timing and magnitude of the MRCP minima are indicative of distinct motor activities, and cortical processes [34, 59, 60]. The MRCP minima can be considered as characteristic features of the movement monitoring potential (MMP) and are hence related to the subject’s intent to correct errors when following the force profile [38,61]. The scaling of the RFD levels with the time between the most negative local minima (Fig. 2(a)) is the most interesting observation on the MRCP features that we have reported here.

The MRCP statistical characteristics demonstrated 75% classification accuracy, highlighting the role of *δ*-band signals in motor tasks. This strong classification accuracy illustrates the potential of MRCP features to map to RFD, confirming the first hypothesis. Even though the full spectrum of EEG from 9 channels (9774 features) can secure the performance of 84% which is higher than the performance of *δ*-band MRCP, however the results support that the small feature set of statistical MRCP characteristics can recover the classification performance and closely follow that of the full spectrum, confirming the second hypothesis. Furthermore, the two-level classification accuracy with the MRCP statistical characteristics was at ~ 90%, suggesting strong discrimination of the behavior based on the MRCP waveform when the Ballistic and Fast trials are compared with the Slow and Medium trials. Our preliminary analysis of applying the STFT only on the *δ*-band showed an overall average accuracy of 67%, underlining the advantage of the MRCP statistical characteristics compared to the STFT feature set of the same signal. The strong classification accuracy of MRCP statistical characteristics (with very low number of features) is aligned with the individual features showing statistical differences between the levels of RFD (Fig. 3(b)); for example, our analysis showed that the Slow task can be differentiated from Ballistic by the signal mean, slope and standard deviation features (*p* < 0.008). The MRCP statistical characteristics are indicative of the complexity of the waveform and also the rate and overall magnitude (and power) of the deflection of the waveform from its baseline. The combination of these statistical characteristics quantitatively describes the changes in the MRCP signal, and, based on the results, can effectively map the signal to the different RFD levels.

All feature sets quantitatively demonstrated that non-adjacent classes of RFD were rarely confused with one another (Figs. 4(b), 5(b), 7(b)). The MRCP statistical characteristics and full-spectrum STFT feature sets achieved strong accuracy thresholds with a simple SVM algorithm. Despite the extensive literature on the use of BCI for decoding spatial/kinematic aspects of motion (mainly in the upper limb [47,62,63]), literature is limited for decoding kinetic aspects of motor intention, specifically the intensity of task conduction. In this regard, in the last decade, research has been conducted to predict two levels of RFD using EEG signals in the lower limb [43, 55]. However, to the best of the knowledge of the authors, there has been no study on discriminating greater than two intensity levels. A higher number of detectable intensity levels means a higher resolution of the BCI in decoding the intensity of intended motion, which is imperative for intuitive implementations. In this paper for the first time, we investigate the information content of MRCP for decoding up to 4 levels of intensity. It should also be noted that recently complex machine learning algorithms, such as deep learning architectures, have been used for processing various biosignals, such as EMG [64, 65], EEG [66, 67], and more specifically neural traces such as MRCP [44, 68]. Using such techniques, often high performance is achieved. However, the high dimensionality of such models creates difficulty in identifying key features and neurophysiologically interpreting the results. Also, complex models often call for a large volume of training datasets to secure the performance which is not always feasible, especially for bio-signals. In this study, we show that using a minimal neurophysiologically-meaningful MRCP feature set can provide adequate information about the kinetic output and successfully classify the response with up to four levels of intensity (which was not reported before). Also, using known feature sets with relatively low dimensionality can help with the interpretability of the results, especially in the case of identifying a motor impairment.

Limitations of this study were that we did not perform source localization on the EEG. The features pertaining to the MRCP deflections before the start of the task (RP1 slope, RP2 slope, and min1) are components of motor preparation and likely to originate from the PMd [28, 29, 37]. The rest of the waveform may represents a mixed activity from the primary motor cortex, SMA, and other neighboring motor cortices [29, 69].

The illustrated relationship between MRCP and RFD has significant potential for application to neurorehabilitation and BCI by providing information about the patient’s intended kinetic parameters (i.e., force and especially RFD), so that the assistance can be tailored to the patient’s intended task speed. We have shown that the accuracy of the classified RFD is superior after, rather than before, the movement onset (Fig. 6, Table 1). The MRCP features after the movement showed the highest effect size (Fig. 3), which explains the superior classification performance. Harnessing the post-movement discriminative power, the MRCP could determine the needed assistive RFD for patients with sensorimotor impairments, in tandem with the intention-detection algorithms [39, 41, 42, 70], improving the quality of neurorehabilitation. In light of the use of the MRCP to trigger stimulation and thus induce neuroplastic changes in the nervous system [29, 71–73], the stimulation could be adjusted based on the classified target RFD level [8, 74]. Finally, the MRCP has been demonstrated as a biomarker of stroke [75], hence novel MRCP features revealed here may prove useful in reinforcing this biomarker to discriminate impairments and motor improvements.

### Disclaimer

This article reflects the views of the authors and should not be construed to represent FDA’s views or policies. The mention of commercial products, their sources, or their use in connection with material reported herein is not to be construed as either an actual or implied endorsement of such products by the Department of Health and Human Services.

## Acknowledgements

This material is based upon work supported by the National Science Foundation (Awards # 2037878, 2229697).

